# Micro- and nanostructural characteristics of rat masseter muscle entheses -Characteristics of masseter muscle entheses-

**DOI:** 10.1101/648105

**Authors:** Keitaro Arakawa, Satoru Matsunaga, Kunihiko Nojima, Takayoshi Nakano, Shinichi Abe, Masao Yoshinari, Kenji Sueishi

## Abstract

The entheses of the masticatory muscles differ slightly from those of the trunk and limb muscles. However, the bones of the skull are subject to various functional pressures, including masticatory force, resulting in a complex relationship between bone structure and muscle function that remains to be fully elucidated. The present study aimed to clarify aspects of masseter muscle-tendon-bone morphological characteristics and local load environment through quantitative analysis of biological apatite (BAp) crystallite alignment and collagen fiber orientation together with histological examination of the entheses.

Result of histological observation, the present findings show that, in the entheses of the masseter muscle in the first molar region, tendon attaches to bone via unmineralized fibrocartilage, while some tendon collagen fibers insert directly into the bone, running parallel to the muscle fibers. Furthermore, BAp crystallites in the same region show uniaxial preferential alignment at an angle that matches the insertion angle of the tendon fibers. Conversely, in the entheses of the masseter muscle in the third molar region, the tendon attaches to the bone via a layer of thickened periosteum and chondrocytes. As in the first molar region, the results of bone quality analysis in the third molar region showed BAp crystallite alignment parallel to the orientation of the tendon fibers. This indicates that the local mechanical environment generates differences in enthesis morphology.

The present study showed a greater degree of uniaxial BAp crystallite alignment in entheses with direct insertion rather than indirect tendon-bone attachment and the direction of alignment was parallel to the orientation of tendon fibers. These findings suggest that functional pressure from the masseter muscle greatly affects bone quality as well as the morphological characteristics of the enthesis, specifically causing micro- and nanostructural anisotropy in the direction of resistance to the applied pressure.

## 1. Introduction

The tendons of the trunk and limb muscles attach to bones via entheses [1–3]. These attachment sites are broadly categorized as either fibrous entheses, composed of perforating mineralized collagen fibers, or fibrocartilaginous entheses, comprising a multitissue interface involving the following four tissues of tendon, unmineralized fibrocartilage, mineralized fibrocartilage, and bone [4]. The composition, structure, and mechanical properties of these multitissue interfaces vary widely, creating spatial gradients that mediate load transfer between soft and hard tissues and minimize stress concentration [5]. Muscle loading is extremely important for healthy enthesis formation and suppression of muscle function greatly diminishes the biomechanical performance of the enthesis [6].

Histological examination by Hems et al. revealed that the entheses of the masticatory muscles differ slightly from those of the trunk and limb muscles [7]. Specifically, the masticatory muscles contain three types of enthesis, including sites of direct tendon insertion into the bone. The authors concluded that these different types contribute to the unique biomechanical function of the masticatory muscles, enabling them to work as an “angle and stretching brake”. However, the bones of the skull are subject to various functional pressures, including masticatory force, resulting in a complex relationship between bone structure and muscle function that remains to be fully elucidated. Clarification of the relationships between the micro- and nanostructural characteristics of the muscles, tendons, and bones in the maxillofacial area and the mechanical environment is required [8, 9].

The relevance of bone quality in addition to bone density with regard to bone strength was proposed National Institutes of Health Consensus Development Conference in 2000. Since then, studies on the relationship between bone structural characteristics and bone strength have focused on bone quality factors [10]. Collagen fibers and biological apatite (BAp) crystallites have been identified as dominant bone quality factors that are resistant to tensile and compressive stress, respectively, on bone tissue [11, 12].

Biological apatite is a hexagonal, ionic crystal that has a highly anisotropic nanostructure with preferential alignment along the c axis in the loading direction [13]. Using microbeam X-ray diffraction analysis, Nakano et al. quantitatively analyzed BAp crystallite alignment in animal trunk and limb bones, demonstrating a high correlation between mechanical stress and BAp crystallite alignment [14, 15]. With regard to collagen fibers as a bone quality factor, ongoing research by Vashishth et al. has shown that collagen crosslinks are a factor in age-related reductions in bone quality [16]. Meanwhile, Kawagoe et al. reported a relationship between bone strength and orientational anisotropy of collagen fibers for the masticatory muscles [17]. Quantitative analysis of the jaw bone, particularly at entheses, should enable accurate prediction of the effects of the masseter muscles on the load environment of the mandible.

The present study aimed to clarify aspects of masseter muscle-tendon-bone morphological characteristics and local load environment through quantitative analysis of BAp crystallite alignment and collagen fiber orientation together with histological examination of the entheses.

## 2. Materials and Methods

### 2.1 Samples

The present study was approved by the Ethics Committee of Tokyo Dental College (Ethics Application No. 282807). Samples were prepared from the skulls of five 24-week-old male Wistar rats euthanized after deep anesthesia with ethyl ether.

### 2.2 Tissue slice preparation

To obtain suitable samples for bone quality analysis, the left skull was embedded in autopolymerizing acrylic resin and sagittally sectioned using a saw microtome (SP1600; Leica, Wetzlar, Germany) with a blade width of 300 μm. Samples were then sanded using wet/dry sandpaper of increasing grit (400, 800, and 1200) to prepare thin, 200 μm slices. The right skull was fixed in 4% paraformaldehyde phosphate buffer solution and demineralized in 10% ethylenediaminetetraacetic acid (EDTA) for 4 weeks. Using standard methods, samples were embedded in paraffin embedding and sliced about 5µm thick in the coronal plane to enable observation of the masseter muscle entheses. Masson’s trichrome staining were performed to observe the structural morphology of masseter muscle entheses in the first and third molar regions. And Toluidine blue staining were used to make the acidic mucus polysaccharide present in the cartilage metachromatic.

### 2.3 Second harmonic generation (SHG) imaging

SHG images were acquired using a multiphoton confocal microscopy system (A1R+MP, Nikon, Japan) with an excitation laser (Mai Tai eHP, wavelengths: 690-1040 nm; repetition rate: 80 MHz; pulse width: 70 fs; Spectra-Physics, CA, US) and a water-immersion objective lens (CFI75 Apo 25×W MP, numerical aperture: 1.1; Nikon, Tokyo, Japan). The excitation wavelength for the observation of collagen fibers was 880 nm. Image acquisition, processing for orthogonal views and cropping were performed using NIS-Elements version 4.0 (Nikon). Brightness and contrast of some images were adjusted using look-up tables (LUTs) of this software by the same parameters among relevant images to facilitate visibility.

### 2.4 Micro-computed tomography (micro-CT) imaging

Samples were examined using micro-computed tomography (micro-CT; HMX225 Actis4; Tesco, Tokyo, Japan) under the following imaging conditions: tube voltage, 140 kV; tube current, 100 µA; matrix size, 512×512; magnification, ×2.5; slice width, 50 µm; and slice pitch, 50 µm. Three-dimensional reconstruction was performed using TRI/3D-BON software (RATOC System Engineering, Tokyo, Japan).

### 2.5 BAp crystallite alignment

Quantitative analysis of BAp crystallite alignment was conducted using an optical curved imaging plate (IP) X-ray diffraction system (XRD; D/MAX PAPID II-CMF; Rigaku, Tokyo, Japan). Measurements were performed in reflection and transmission modes with Cu-Kα as the radiation source at a tube voltage of 40 kV and tube current of 30 mA. Reference axes were established in X axis, Y axis, and Z axis for each sample (Fig. 1) [18, 19]. Regions of interest in the mandible comprised masseter muscle entheses (Fig. 2). The radiation site was determined using the light microscope of the XRD system (magnification, ×0.6-4.8), then an incident beam (diameter, 50 μm) was applied. Using reflection mode in the X-axis direction and transmission mode in the Y-axis and Z-axis directions, the diffracted X-ray beam was detected using a curved IP based on the conditions described by Nakano et al. [20]. The diffracted X-ray beam was detected as a diffraction ring on the IP. Using 2-dimensional data-processing software (Rigaku), X-ray diffraction intensity ratios were calculated for the two diffraction peaks corresponding to planes 002 and 310.

**Fig 1.**
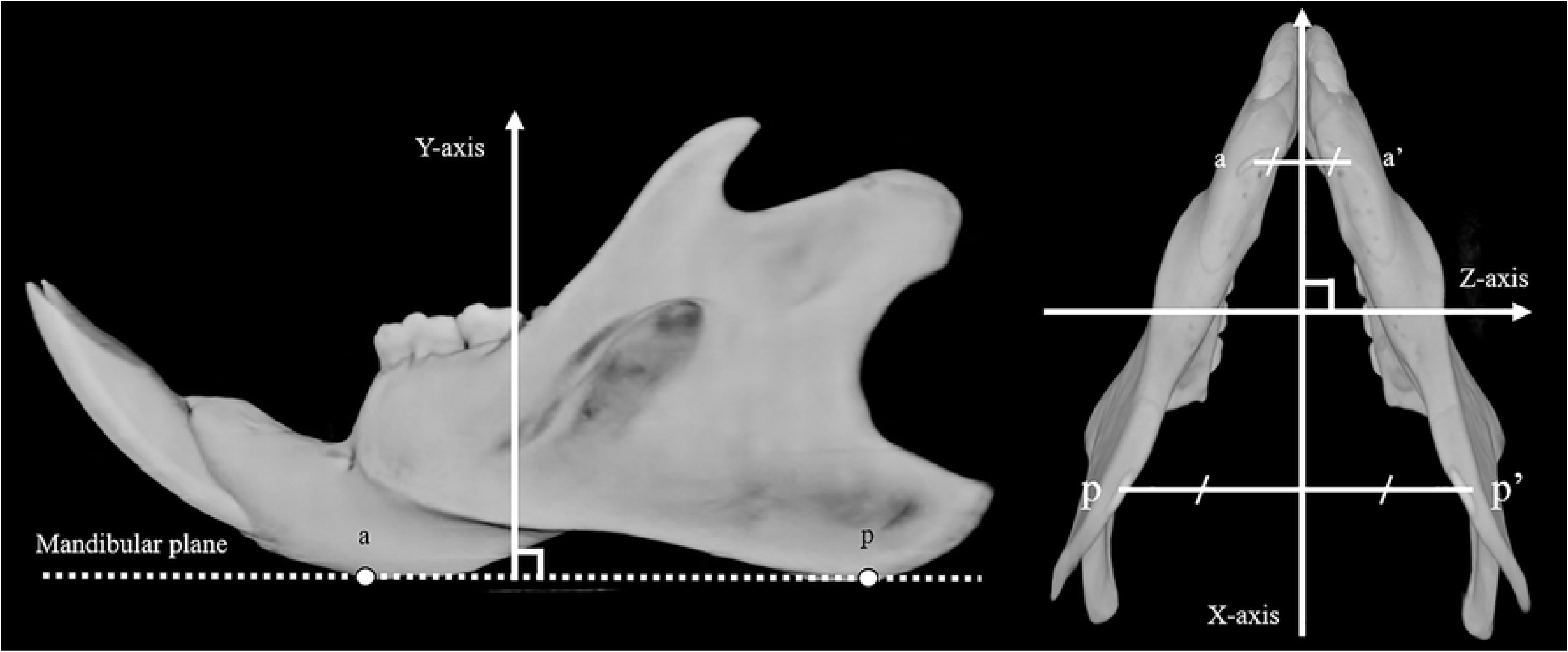
Reference points, plane and axes: point a the lowest point in anterior thickening area of mandible. Point p the lowest point in posterior thickening area of mandible; Mandibular plane passing through a-a’ and p-p’ lines; X-axis passing through the mid-point of a-a’ and p-p’; Y-axis the vertical axis against the mandibular plane; Z-axis the vertical axis against the X-Y plane.

**Fig 2.**
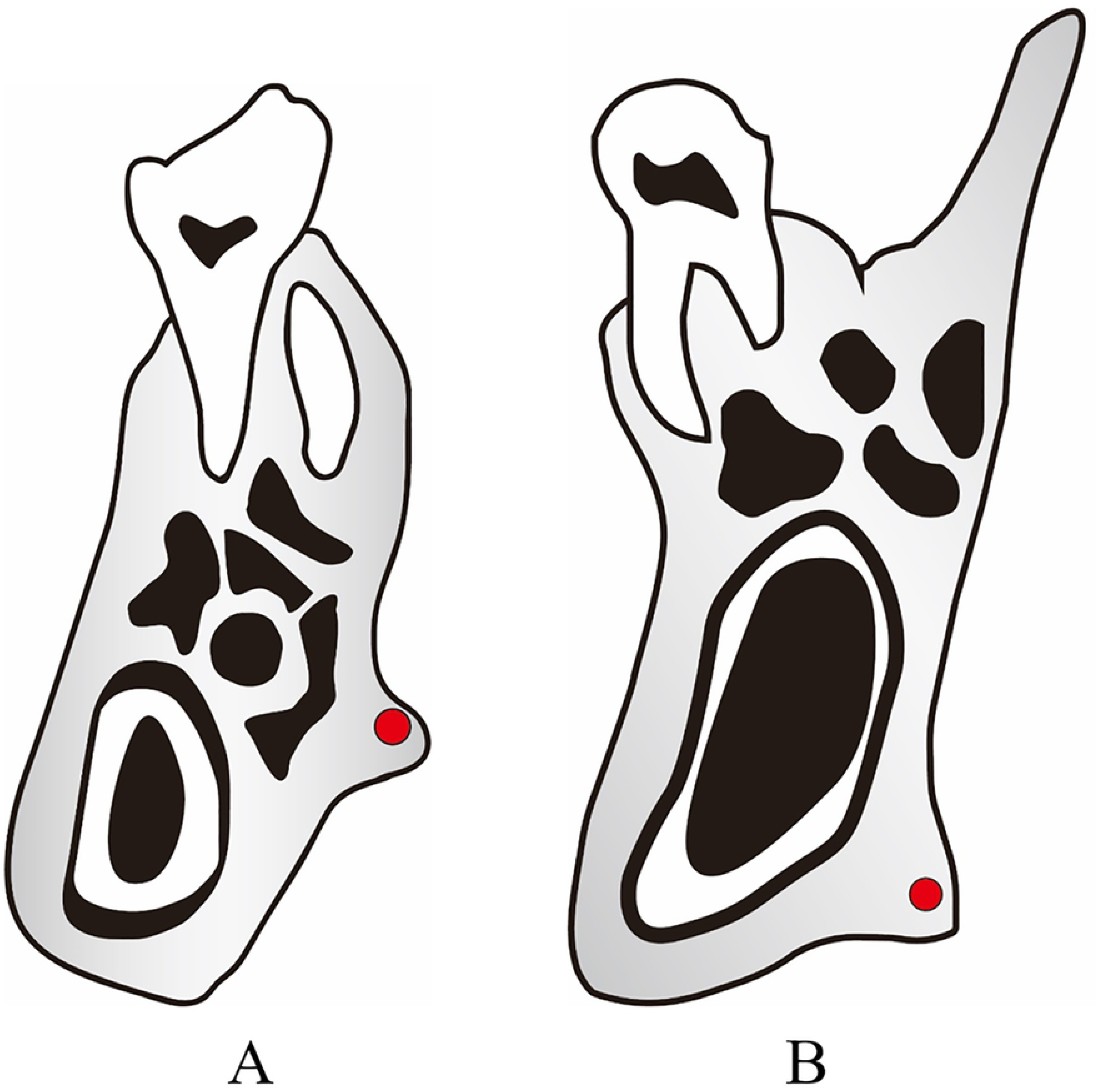
Measurement points. (A) First molar region. (B) Third molar region.

### 2.6 Statistical analysis

Mean values for the five samples for each measurement point were calculated and compared using Tukey’s multiple comparison test. Significance was set at *P*<0.05.

## 3. Results

### 3.1 Histological observation of entheses

The results of Masson’s trichrome staining and toluidine blue staining are showed of a coronal section in the first molar region of 24-week-old Wistar rat skulls (Fig. 3). Thick masseter muscle tendons with fibers largely grouped into bundles could be seen directly integrating into the buccal cortical bone. In the enthesis, the periosteum was fragmented and aggregation of chondrocytes was observed at the tendon-bone interface (Fig. 3B, C,D,E). The results of Masson’s trichrome staining and toluidine blue staining are showed of a coronal section in the third molar region of 24-week-old Wistar rat skulls (Fig. 4).

**Fig 3.**
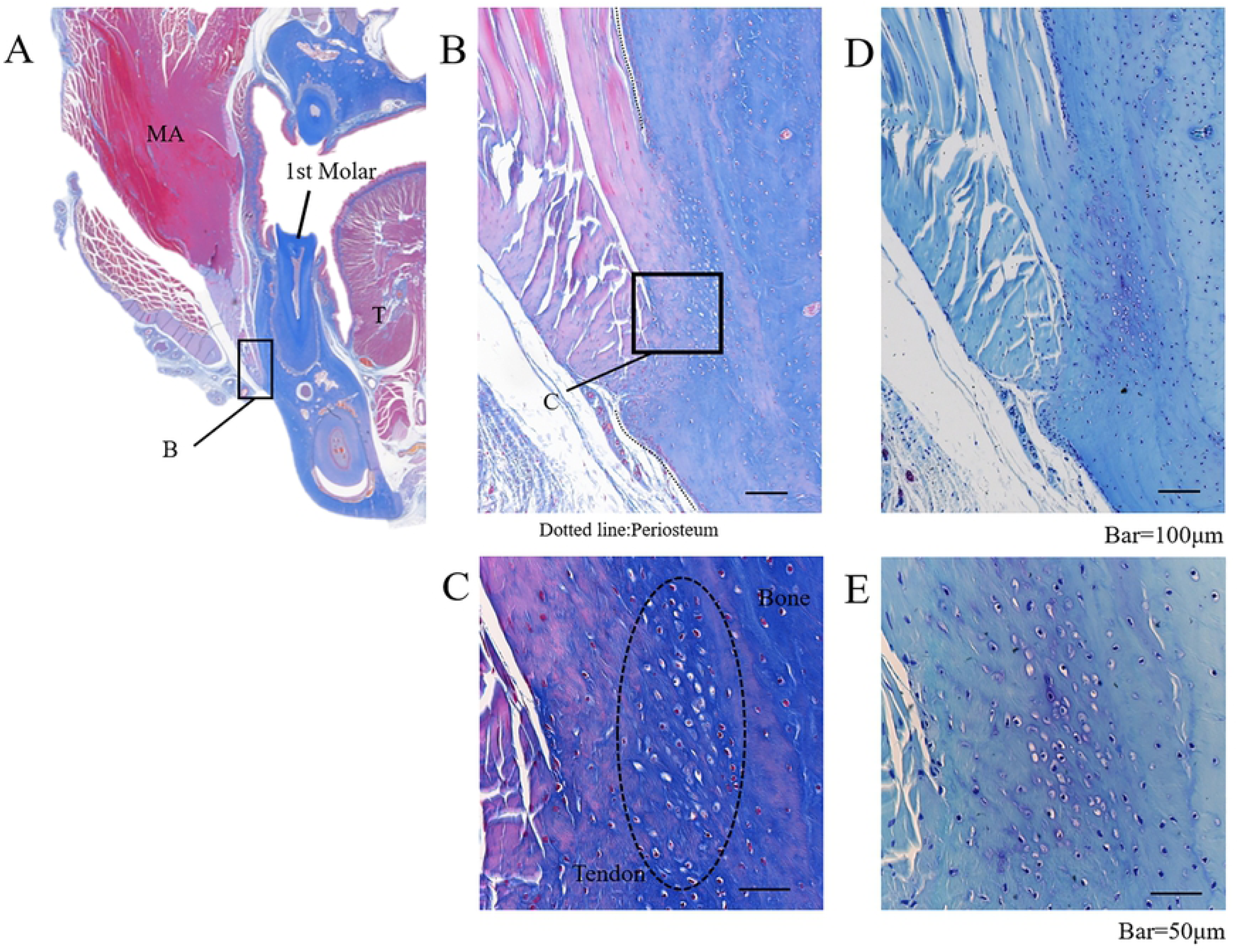
Masson’s trichrome and toluidine blue staining of masseter muscle enthesis in the first molar region. MA: masseter muscle. T: tongue. 1st Molar: first molar region.

**Fig 4.**
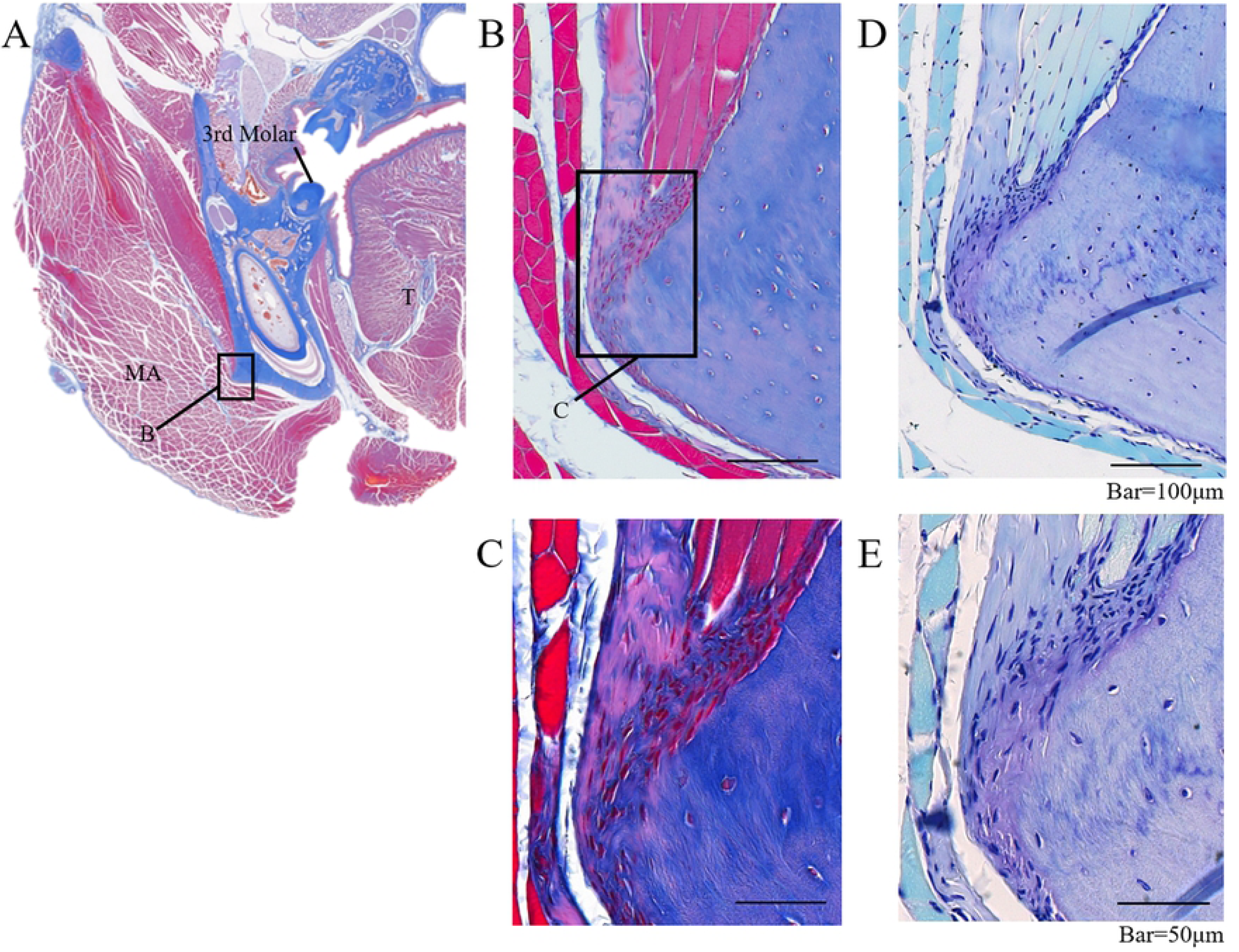
Masson’s trichrome and toluidine blue staining of masseter muscle enthesis in the third molar region. MA: masseter muscle. T: tongue. 3rd Molar: third molar region.

In the masseter muscle enthesis, muscle fibers could be seen covering the area from the lateral aspect to the base of the mandible, while thin tendons ran toward the cortical bone protuberance on the buccal side of the mandibular body (Fig. 4A). The tendon was attached to the periosteum without rupturing (Fig. 4B,D) and thickened chondrocytes were observed on the bone surface in the enthesis (Fig. 4C,E).

### 3.2 Orientational anisotropy of collagen fibers

SHG images from the first molar region are showed in Figure 5. The masseter muscle tendon in the first molar region comprised thick collagen fibers extending through the enthesis, some of which penetrated the cortical bone. As in the first molar region, collagen fibers in the third molar region ran toward the bone. However, these fibers were interrupted at the thickened periosteum. In the vicinity of the enthesis, many collagen fibers inside the bone were observed running parallel to the orientation of the tendon (Fig. 6). In the first and third molar regions, tendon fibers ran tangentially to the bone at 31.1° [standard deviation (SD), 4.6°] and 40.4°[SD, 3.5°], respectively (Fig. 7).

**Fig 5.**
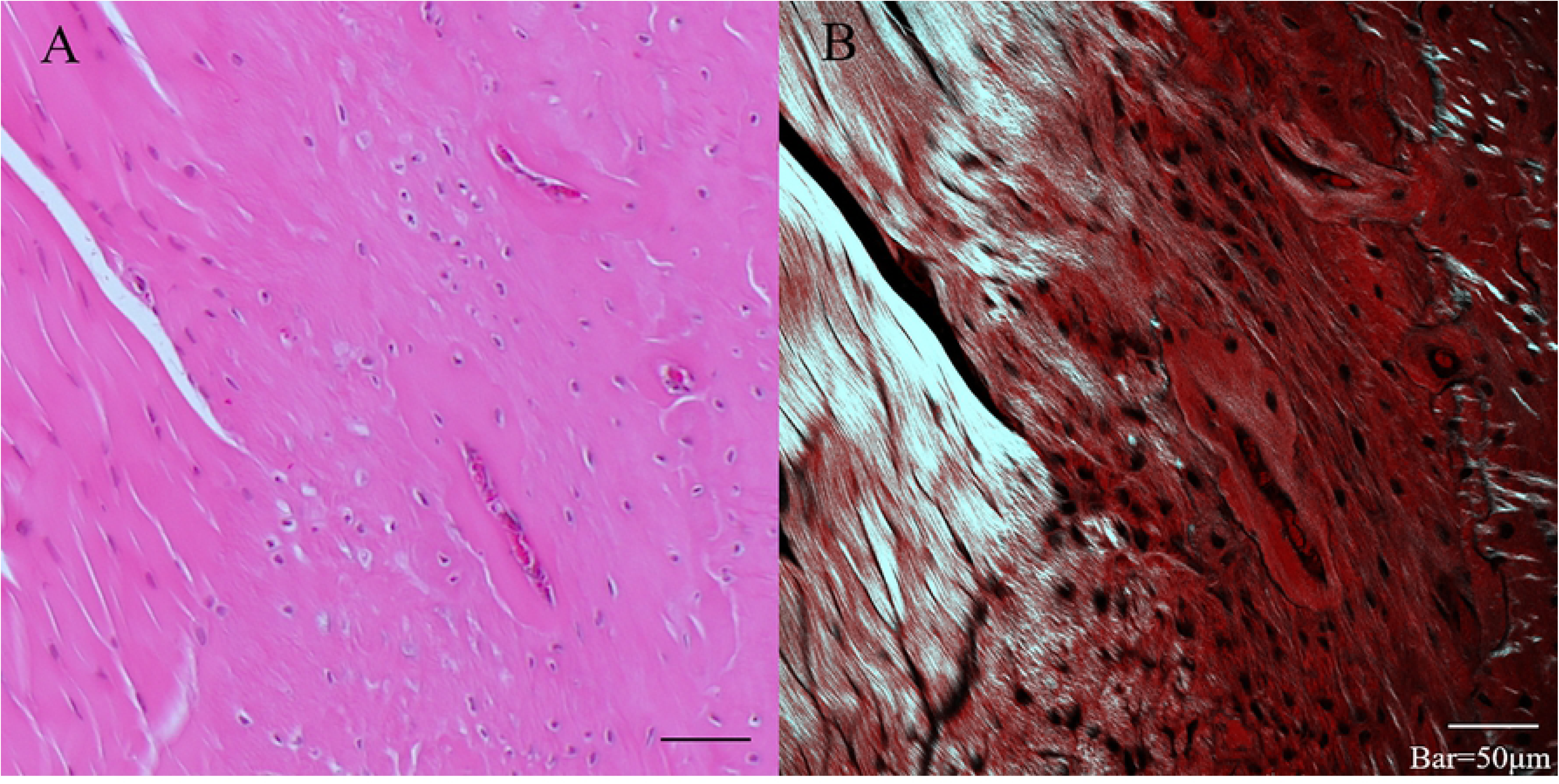
Coronal section in first molar region. (A) Hematoxylin and eosin staining. (B) Second harmonic generation imaging. The masseter muscle tendon in the first molar region comprised thick collagen fibers extending through the enthesis, some of which penetrated the cortical bone.

**Fig 6.**
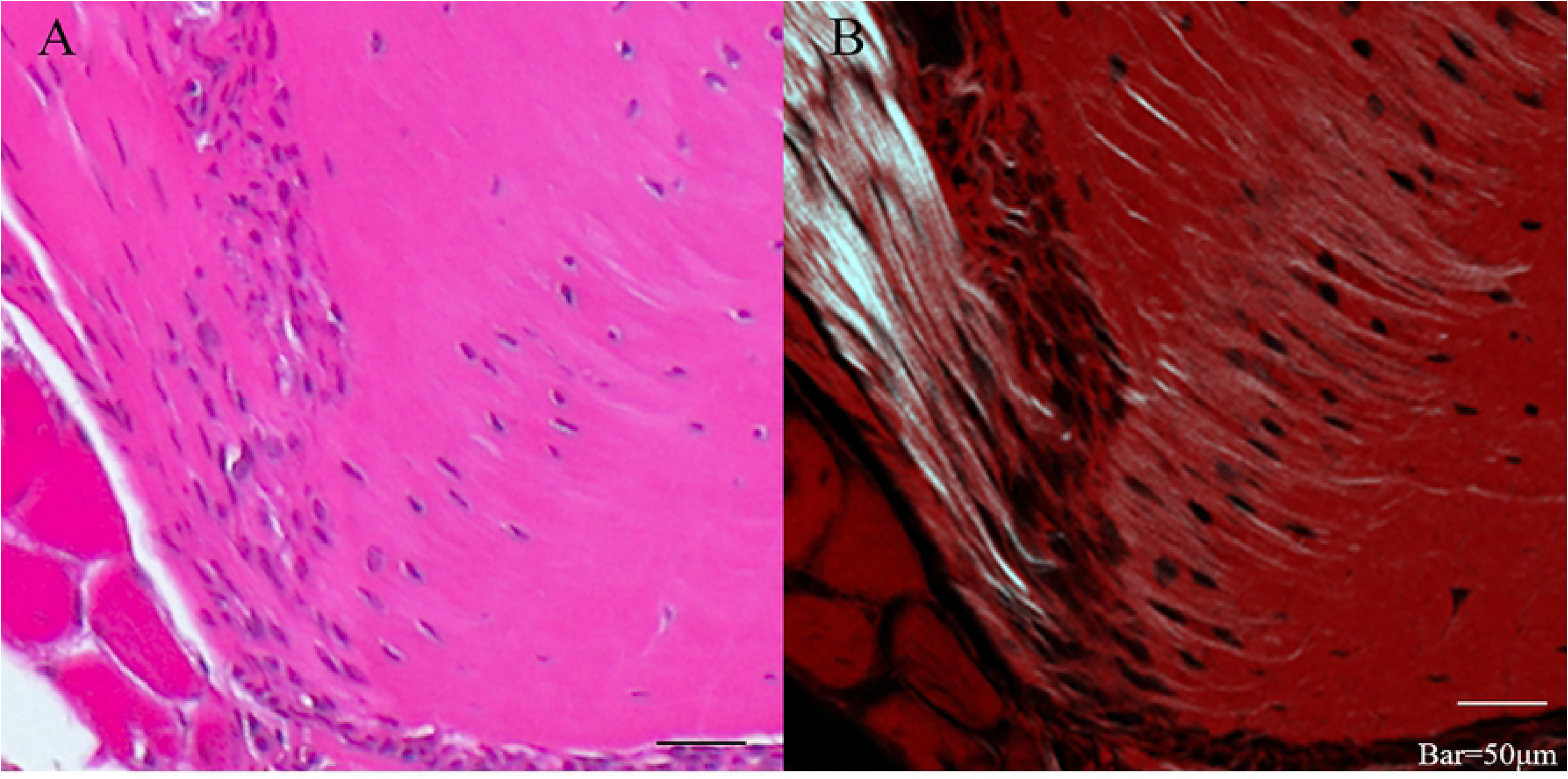
Coronal section in third molar region. (A) Hematoxylin and eosin staining. (B) Second harmonic generation imaging. In the vicinity of the enthesis, many collagen fibers inside the bone were observed running parallel to the orientation of the tendon.

**Fig 7.**
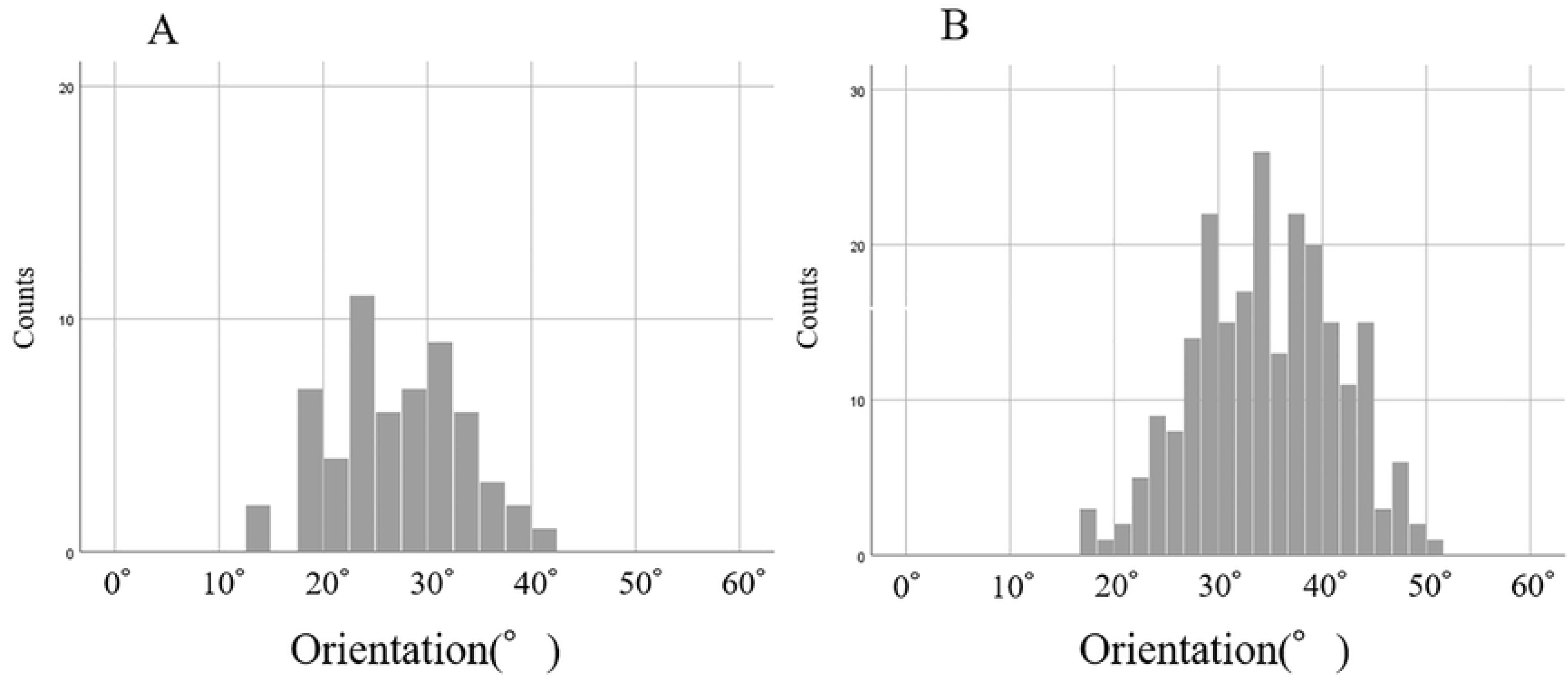
Orientation of tendon fibers in relation to bone. (A) First molar region. (B) Third molar region.

### 3.3 BAp crystallite alignment

The angles of preferential alignment of BAp crystals are showed in an enthesis in relation to tooth axis in the first and third molar regions (28° [SD, 10.95°]) and 36° [SD, 8.94°], respectively). X-ray diffraction intensity ratios calculated for the three reference axes for quantitative analysis are showed in Figure 8. The intensity ratios for hydroxyapatite powder were 1.4 and 5.6 in reflection and transmission modes, respectively. In both the first and third molar regions, strong uniaxial preferential alignment was noted in the Y-axis direction in the masseter muscle entheses. Furthermore, X-ray diffraction intensity ratios in the Y-axis direction were significantly higher in the first compared to the third molar region.

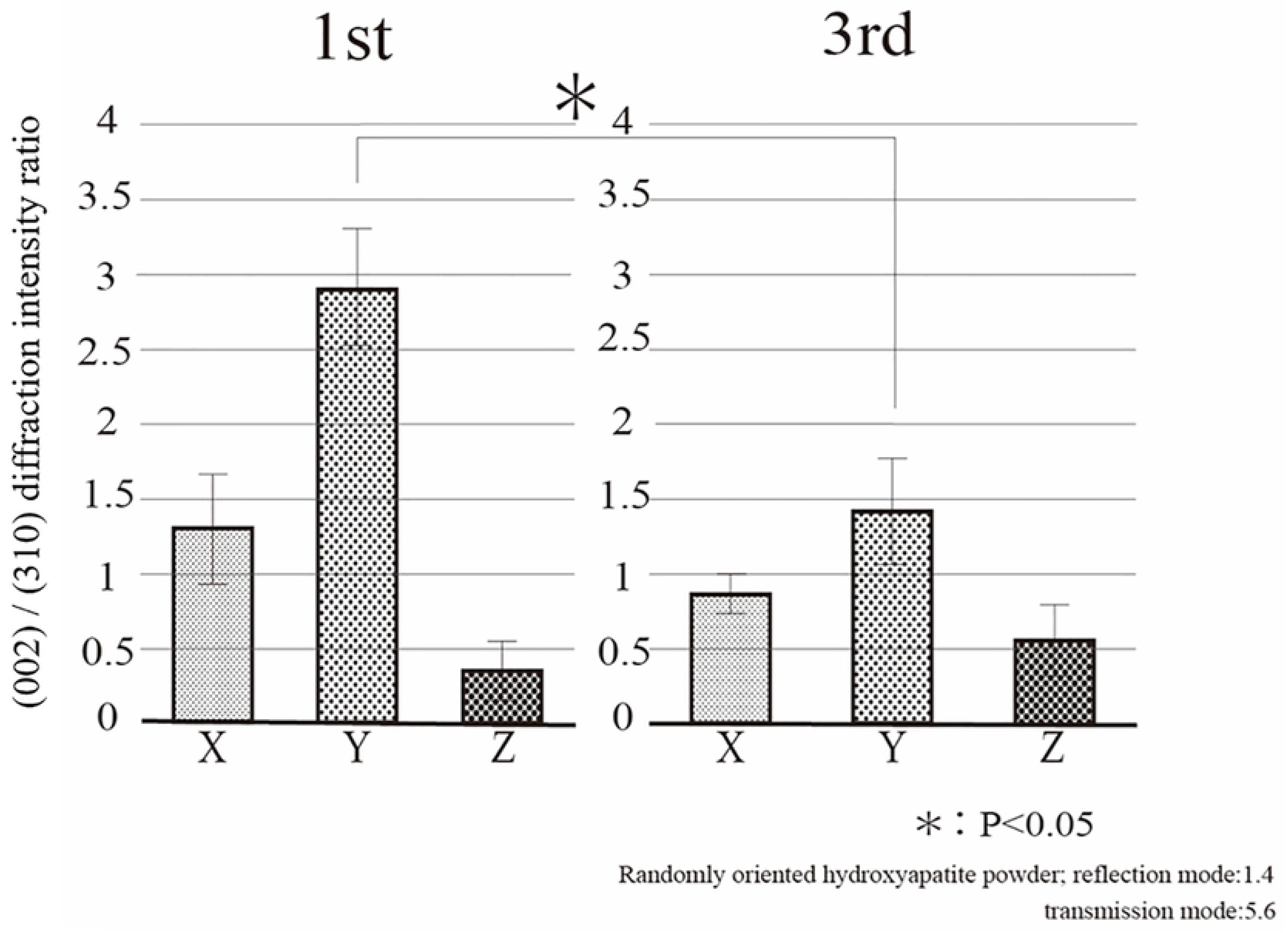

## 4. Discussion

According to Huang et al., as the superior digital flexor tendon develops before and then joins with the subsequently formed digital bone, the related entheses penetrate the periosteum [21]. Conversely, the entheses of many of the muscles in the trunk and limbs do not involve the periosteum, suggesting that these attachments are already formed at the stage of periosteal development [22]. The present findings also show that, in the entheses of the masseter muscle in the first molar region, tendon attaches to bone via unmineralized fibrocartilage, while some tendon collagen fibers insert directly into the bone, running parallel to the muscle fibers. Furthermore, BAp crystallites in the same region show uniaxial preferential alignment at an angle that matches the insertion angle of the tendon fibers. This suggests that both the anisotropy of the collagen fibers and BAp crystallite alignment confer high resistance in the direction of the masseter muscle tendon. As the masseter muscle in the first molar region is directly attached to the bone via the tendon, these structural characteristics may optimize bone quality to enable the high load generated by muscle contraction to be efficiently transmitted from tendon to bone [23].

Conversely, in the entheses of the masseter muscle in the third molar region, the tendon attaches to the bone via a layer of thickened periosteum and chondrocytes. Muscles with entheses that indirectly attach to the bone via the periosteum do not produce large functional pressures; rather, they are responsible for precise movements [24]. As in the first molar region, the results of bone quality analysis in the third molar region showed BAp crystallite alignment parallel to the orientation of the tendon fibers. However, the intensity ratio values were significantly lower. This indicates that the local mechanical environment generates differences in enthesis morphology.

Matsumoto et al. analyzed bone quality in human jaw bones and found uniaxial preferential alignment of BAp crystallites and high bone strength in the tooth axis direction in specific alveolar bone [25]. Meanwhile, Nakano et al. demonstrated a strong positive correlation between bone quality factors and bone strength, indicating that the mechanical environment of entheses determines the orientation of the collagen fibers, which is linked to the preferential alignment of BAp crystallites. Changing the amount and direction of functional pressure from the muscles may thus affect not only bone density, but also bone density.

## 5. Conclusion

In the entheses of rat masseter muscle, some tendons attach to the bone directly and others attach indirectly via the periosteum. The present study showed a greater degree of uniaxial BAp crystallite alignment in entheses with direct insertion rather than indirect tendon-bone attachment and the direction of alignment was parallel to the orientation of tendon fibers. These findings suggest that functional pressure from the masseter muscle greatly affects bone quality as well as the morphological characteristics of the enthesis, specifically causing micro- and nanostructural anisotropy in the direction of resistance to the applied pressure.

## 6. Acknowledgement

This research was supported by a Grant-in-Aid for Scientific Research (C) (17K11808 and 25463055) from the Japan Society for the Promotion of Science. This study was supported by a research grant under the Private University Research Branding Project from the Ministry of Education, Culture, Sports, Science, and Technology of Japan (2018).

## 7. Conflict of Interest

No potential conflicts of interest to disclose.

